# Transpiration and water use efficiency of sorghum canopies have a large genetic variability and are positively related under naturally high evaporative demand

**DOI:** 10.1101/2022.09.21.508841

**Authors:** Raphaël Pilloni, Kakkera Aparna, Zineb El Ghazzal, Soumyashree Kar, A Ashok Kumar, Amir Hajjarpoor, Pablo Affortit, William Ribière, Jana Kholova, François Tardieu, Vincent Vadez

**Affiliations:** Université de Montpellier, IRD, DIADE, Montpellier, France; International Crop Research Institute in Semi-Arid Tropics, Hyderabad, India; Iowa State University, Ames, Iowa, USA; Université de Montpellier, INRAE. Laboratoire d’Ecophysiologie des Plantes sous Stress Environnementaux, Montpellier France; Laboratoire Mixte International LAPSE, Dakar, Sénégal

**Keywords:** Sorghum, canopy density, water use efficiency, vapor pressure deficit, evapotranspiration, Transpiration, genetic variability, light penetration

## Abstract

Indoor experiments with individual plants often show that transpiration rate is restricted under high vapor pressure deficit (VPD), resulting in a plateau of transpiration that increases water use efficiency (WUE) of some genotypes. We tested this hypothesis outdoors during dry or rainy seasons of India and Senegal, based on the response of the transpiration of canopy-grown sorghum plants to the reference evapotranspiration that takes both light and VPD into account. This response showed no plateau at high evaporative demand in 47 genotypes, but a large genetic variability was observed for the slope of the relationship over the whole range of evaporative demand. Unexpectedly, this slope was genetically correlated with WUE in two experiments with high evaporative demand: genotypes that most transpired had the highest WUE. Conversely, a negative correlation was observed under low evaporative demand. Genotypes with high WUE and response to evaporative demand were also those allowing maximum light penetration into the canopy. We suggest that this caused the observed high WUE of these genotypes because leaves within the canopy had sufficient light for photosynthesis whereas we observed a lower VPD in the canopy than in open air when leaf area index reached 2.5-3, thereby decreasing transpiration.

**Highlights:** The transpiration response to evaporative demand was genetically variable and correlated to WUE: genotypes that most transpired had highest light penetration towards leaves subjected to lower VPD than in air.

## Introduction

Water use efficiency (WUE) tends to decrease with vapor pressure deficit (VPD) because transpiration, but not photosynthesis, increases with VPD (Sinclair, Tanner and Bennett, 1984; Condon *et al*., 2002, 2004). A mechanism used by plants to avoid such a decrease is stomatal closure at high VPD, which affects transpiration more than photosynthesis because of the non-linear relation between these two variables (Farquhar and Sharkey, 1982). Hence, restricting transpiration under high VPD theoretically increases WUE, especially during hottest hours of the day, provided that no confounding effects occur. Indeed, genotypes that most close stomata during the afternoon were reported to have highest WUE (Sinclair et al., 2005, Vadez et al., 2014). These analyses have triggered research on the genetic variability of the transpiration restriction under high VPD (Fletcher et al., 2008; Vadez et al., 2014). Genetic variation in the transpiration response to increasing VPD was observed in several species such as soybean (Fletcher, Sinclair and Hartwell, 2008), wheat (Schoppach and Sadok, 2012), pearl millet (Kholová *et al*., 2010; Choudhary *et al*., 2020), maize (Gholipoor *et al*., 2013; Choudhary *et al*., 2020), sorghum (Gholipoor, Sinclair and Prasad, 2012; Choudhary *et al*., 2020) and rice (Affortit *et al*., 2022). A genetic variability for WUE was also described in sorghum (Donatelli et al., 1992, V. Vadez et al., 2011; Xin et al., 2009), and was interpreted as a consequence of the plant ability to restrict transpiration in response to evaporative demand, thereby generating a plateau of transpiration rate at high VPDs (Sinclair, Hammer and Van Oosterom, 2005; Hatfield and Dold, 2019).

Intriguingly, plateaus of transpiration under increasingly high VPD were essentially observed in growth chamber experiments, with constant light at intensities usually lower than 500 μmol m^-2^ s^-1^. Yet, light intensity outdoors is often much higher, e.g. in sahelian conditions with light intensity higher than 2000 μmol m^-2^ s^-1^ (Adeniji, Adeniji and Ojeikere, 2020), and fluctuates in the course of the day. Results for transpiration under naturally occurring light and VPD did show a trend for a plateau (e.g. Fig.9B in Vadez et al., 2015), although not in all days. Smoothened transpiration profiles also showed a transpiration restriction in some wild chickpea cultivars (Kar et al., 2020), although not as clear as in the theoretical figure 1 of Sinclair et al., (2005). A specific difficulty occurs when examining the relationship between transpiration and evaporative demand in naturally fluctuating light conditions. Transpiration is indeed driven by light intensity and wind in addition to VPD, resulting in the reference evapotranspiration (ETref) as computed with the Penman-Monteith equation (Zotarelli *et al*., 2014). Because VPD and light intensity are often correlated during a day and between sites, several field studies considered VPD only as the driving variable of the evaporative demand (e.g. Schoppach et al., 2017). This assumption may be acceptable in regions in which no massive light variations occur between days and fields during the crop cycle e.g. during dry seasons in India (Kar et al., 2020), but less so in the opposite case. Hence, the genetic variability of the response of transpiration to the evaporative demand needs to be analyzed in natural conditions with light variations, in addition to variations of VPD. We developed a method for that and applied it to two panels of genotypes in three environmental scenarios, namely the dry and wet season in the drylands of India and a dry season in the Sahel of Senegal.

**Figure 1:**
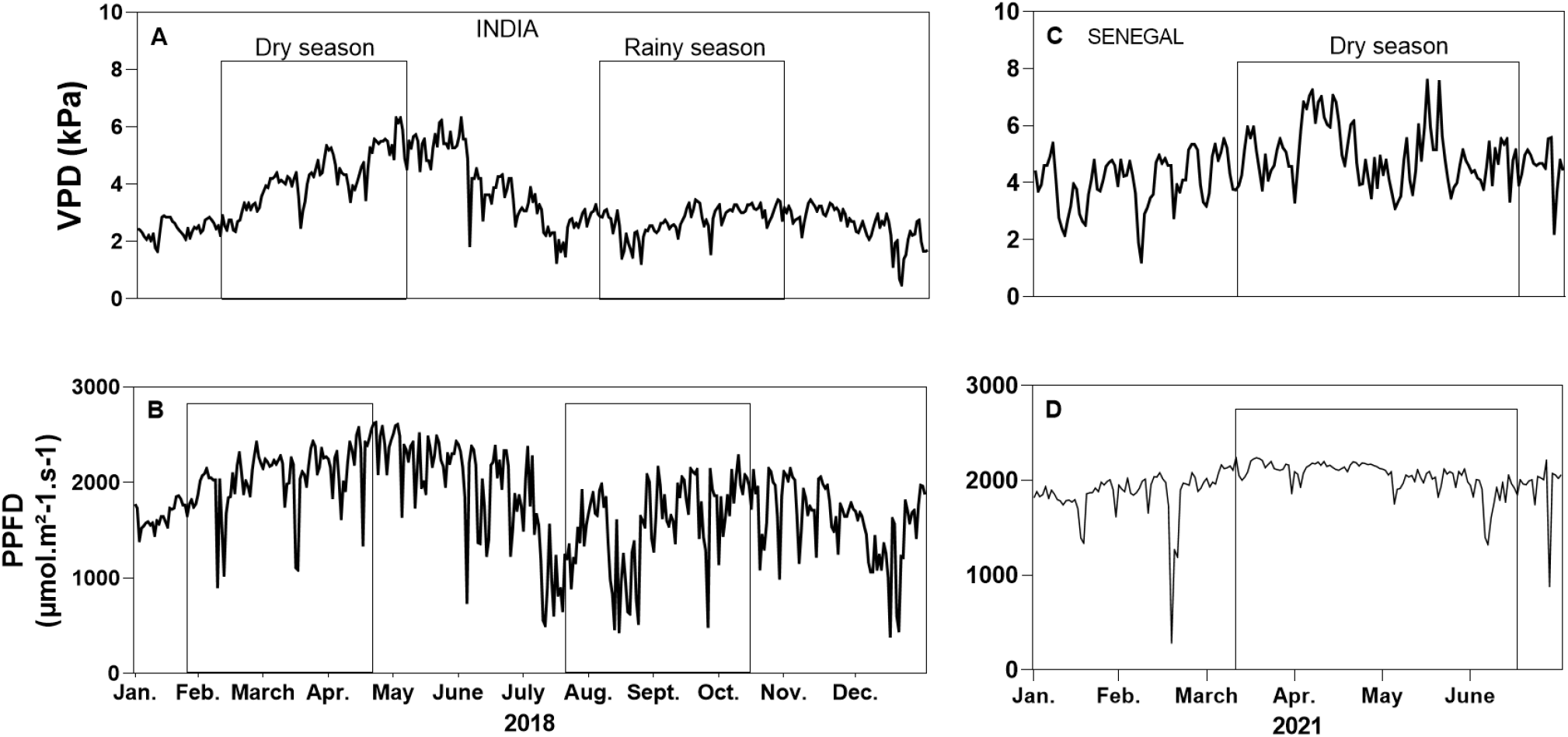
Daily vapor pressure deficit (VPD) and Photosynthetically active Photon Flux Density (PPFD) in experiments in India (ICRISAT meteorological station in 2018, A and B) and Senegal (CNRA Bambey station for the VPD, (C) and Niakhar meteorological station for PPFD in 2021, (D)). Empty boxes correspond to the periods of lysimetric measurements.

Another difficulty for translating the analyses of the genetic variability of transpiration rate from growth chamber to field is associated to the vertical variations of light and VPD. Plants in growth chambers have all leaves exposed to similar light and VPD (predominance of diffuse light, low competition for light and similar VPD at all altitudes in the growth chamber). This differs from plants grown in canopies, where the leaves inside the canopy are expected to be subjected to a lower VPD than the upper leaves because of canopy transpiration, and to a lower light because of self-shading. Provided they receive sufficient light, leaves inside the canopy, with partly closed stomata because of lower light and facing a milder VPD than the upper leaves, may have a higher WUE than upper leaves. The proportion of incident light that reaches the lower parts of the canopy, an essential feature under this hypothesis, has a clear genetic variability (Yin and Struik, 2015) related to plant architecture (Niinemets, 2010), plant density (Song, Zhang and Zhu, 2013) and the vertical distribution of leaf area (Perez *et al*., 2019).

We therefore aimed at evaluating to what extent the genetic variability of the transpiration response to evaporative demand and that of WUE, demonstrated in indoor experiments, is also appreciable under natural conditions for two panels of sorghum genotypes grown under high light and high or low VPD in Senegal and India. For that, we examined the response of each studied genotype to evaporative demand, calculated via a reference evapotranspiration (ET_ref_) taking both VPD and light into account, rather than via VPD alone as in previous studies (Fletcher, Sinclair and Hartwell, 2008; Kholová *et al*., 2010). In order to better understand this genetic variability, we tested if the VPD inside the canopy appreciably differed from that in the air in canopies with different leaf area indices, which were obtained by setting up different plant density treatments, over several days and at different times of the day. We finally explored to what extent WUE differences were linked to light penetration inside the canopy.

## Material and methods

### Genetic material, experimental design and growth conditions

A panel of 20 elite hybrids of sorghum provided by seed companies was used in experiments in India (panel A), and a panel of 27 lines from the germplasm collection of ICRISAT was used in all other experiments (panel B). The field experiments in Senegal involved the whole panel B, while sub samples of nine and two lines of the latter panel were analyzed in the indoor lysimetric platform and greenhouse experiments, respectively (see below).

A first set of experiments was carried out in the field lysimetric platform (LysiField) of the ICRISAT station (Hyderabad, India, 17°30 N; 78°16 E; elevation 549 m) during the dry (February to May) and rainy (August to October) seasons of 2018, which largely differed for VPD (4.7 vs 2.8 kPa in average) and light intensity (2217 vs 1545 μmol m^-2^ s^-1^), (Fig. 1). The platform consisted of large pits in which PVC tubes (20 cm diameter,1.20 m length) were arranged side by side and filled with alfisol. A S-type load cell system set up and a block-chain pulley allowed the tubes to be lifted and weighed every 7^th^-10^th^ day. Direct soil evaporation was prevented via beads located on the soil. Each replicate consisted of a set of 4 tubes arranged in square, all tubes carrying the same genotype. Four replicates were used in both experiments, organized as a complete randomized block design with density as the main factor and genotypes randomized. In the high density treatments, each tube carried one plant, vs one tube out of two in the low density treatments leading to plant densities of 20 and 10 plants m^-2^.

A second experiment took place in the field lysimetric facility of the ISRA/CNRA station (Bambey, Senegal, 14° 41 N; 16° 27 W, elevation 20 m) during the 2021 dry season (March to June 2021) characterized by high VPD and light (4.9 kPa and 2068 μmol m^-2^ s^-1^ on average) (Fig 1). The platform had the same characteristic as the Hyderabad platform with small differences: tubes were 25 cm in diameter and 1.50 m long, they were filled with a sandy soil and soil evaporation was prevented via gravels located on the soil. The experimental design was the same as above, except that high and low plant densities were 16 and 8 plants m^-2^, respectively.

Two field experiments were carried out near the lysimeter experiments in India and Senegal, with panels A and B, respectively, with two plant densities. The field experiment in Senegal was carried out at the CNRA Bambey station. Microplots (2 m wide, 4 m long) were sown with one genotype each, with three reps per genotype. The low density plots harbored 11 plant m^-2^ consisting in 4 plant rows, 15 cm between plants within a row, and an inter row spacing of 60 cm. High density (HD) plots harbored 22 plants m^-2^ with an inter row spacing of 30 cm and the same plant distribution otherwise. Before sowing, the field was fertilized with di-ammonium phosphate at a rate of 100kg/ha and top dressing with 100kg/ha urea occurred at four weeks after sowing in both high and low density treatment. The field experiments in India was carried out at the ICRISAT station close to the lysimeter facility. The experimental design was the same as in the Senegal experiment, with the same two plant densities. Before sowing, the field was fertilized with di-ammonium phosphate at a rate of 100kg/ha and top dressing with 100kg/ha urea four weeks after sowing. Daily temperature, hygrometry and solar radiation data were collected on site at the ICRISAT and CNRA Bambey meteorological stations for experiments in India and Senegal, respectively, except for the radiation data in Senegal that was recorded in anther meteorological station located 25 km south from the field.

An experiment was performed in the indoor IRD automated lysimetric platform of Montpellier, France in April-May 2020-21. Light conditions and VPD fluctuated in the greenhouse, with VPD values ranging from 1 kPa at sunrise to 3.5 kPa at 2 pm. 16 L pots were filled with a mix of loamy clay agricultural soil and sand, each containing four plants and organized in mini canopies of 20 plants m^-2^. A replication consisted of nine pots set up on contiguous 2m long and 0.8m wide tables. Three replicates of each genotypes were randomly distributed on the platform. Each pot was a closed mini lysimeter system that avoided any drainage loss during irrigation. Plastic beads covered the soil surface (4-5 cm layer) to avoid water loss due to direct soil evaporation. Pots were weighed every week and immediately irrigated after weighing to precisely compensate the amount of water transpired by plants of the considered pot. Five weeks after emergence, the mini lysimeters were transferred to an automated lysimetric platform recording the weight of each pot every 30 minutes for a 4-day period.

Finally, an experiment was carried out in the greenhouse experiment at Montpellier in April 2020 with a daytime VPD of 3 kPa maintained over the studied period. Plants were grown in four liters squared pots filled with horticultural potting soil. Pots were placed on 1m^2^ tables. The high and low density treatments consisted in 1 m^2^ tables with 24 or 12 pots. Each treatment was replicated three times for each genotype in the glasshouse.

### Evapotranspiration response to the evaporative demand in a lysimeter setup

Evapotranspiration was measured with lysimeters over several days with natural fluctuations of light and VPD. To maximize the range of evapotranspiration, we also varied leaf area index (LAI) by setting up two plant density treatments. Hence, this protocol profoundly differs from that most often used for establishing response curves to VPD, which involves measuring individual plant transpiration over time scale of 30-60 minute per VPD increment in a growth chamber.

Evapotranspiration data, initially measured in kg of water loss per replication (4 tubes), was first converted into mm m^-2^ day^-1^, by dividing raw values by the reciprocal of plant density in each replicate and by the time between two weighing dates. In India, lysimeters were weighed nine and eight times during the rainy and dry seasons, respectively, resulting in 30 evapotranspiration values for each genotype across the two plant densities. In Senegal, lysimeters were weighed 11 times, giving 20 evapotranspiration values for each genotype. In both cases, measurements covered a period of 7-10 days.

The evaporative demand (ET_ref_) during the same periods of time was calculated in two ways. The first method was based on the simulation of evapotranspiration of a reference genotypes by using the APSIM crop simulation model parametrized with a sorghum genotype having a phenology and leaf area similar to those of hybrids of panel A, based on Priestley-Taylor equation (Priesltley and Taylor, 1972). The transpiration of this reference genotype, simulated with unlimited water supply, is akin to an environmental variable, and was considered as a reference evapotranspiration. The second method involved the Penman-Monteith equation corrected by a crop coefficient (Kc) (Zotarelli *et al*., 2014) which takes into account the change with time of LAI, ranging from 0.4 to 0.85 according to plant stages (Piccinni *et al*., 2009). For both methods, we used as inputs the daily temperature, hygrometry, wind and solar radiation data collected on site at the ICRISAT and CNRA Bambey meteorological stations (Solar radiation data was collected at the Niakhar meteorological station located 25km south of the CNRA statin). Daily ET_ref_ values obtained with either method were averaged over the respective measurement periods and plotted against measured evapotranspiration values for the same perios. Only the second method was used for the Senegal experiment because no closely related genotype was parameterized in APSIM. ET_ref_ data from both methods were compared in the experiment in India. Leaf area index (LAI) was measured 33 and 40 days after sowing in the field directly adjacent to the lysimetric platform by using a 1-meter long ceptometer (AccuPAR LP-80). Four measurement were taken in each replication for the 20 genotypes in both high and low density treatments.

### Transpiration response to the evaporative demand in the glasshouse

In the indoor lysimetric facility of Montpellier, a data analysis pipeline (adapted from Kar et al., 2020) allowed estimation of the time course of transpiration over 24 h. The slope of the time course of transpiration was estimated during the three hours preceding the maximum transpiration, corresponding to the maximum VPD window on a particular day. Because the time course of VPD was similar between days, this slope was an indirect way for comparing genotypes for their transpiration response to evaporative demand.

The slope of the time course of transpiration was estimated during the three hours preceding maximum evapotranspiration, corresponding to the maximum VPD window on a particular day. Because the time course of VPD was similar between days, this slope was an indirect way for comparing genotypes for their transpiration response to the evaporative demand. After 4 days on the load cells, the aboveground biomass of these 6-weeks old plants was harvested, dried in an oven during 72h at 60°C, and used to compute WUE.

### Genotypic variation for water use efficiency in canopy-grown plants

At 13 and 14 weeks after sowing in dry and rainy VPD seasons respectively in India, and at 14 weeks in Senegal, the total above ground biomass was harvested from the lysimeters, dried in an oven during 72h at 60°C and weighted. It was used to compute the water use efficiency (WUE, g of biomass accumulated per kg of water loss through evapotranspiration).

### VPD assessment within canopies

Temperature and relative humidity sensors were positioned in the low and high density plots of the field experiment in Bambey in 2021 (TinyTag ultra 2, TGU-4500, Gemini Datalogger Ltd, Chichester, UK). A daily average of temperature and hygrometry was recorded at the Bambey meteorological station. Air and canopy VPD were calculated based on relative humidity (RH) and air temperature values measured every 30 minutes, either 2 meters above the canopy for air VPD (i.e. from the meteorological station located 50m from the field) or within the canopy at mid height of the plants. Every week, the position of the sensors was adjusted according the height of the plants to be continuously placed at mid height of the canopies. Recording of data started at four weeks after emergence.

Similar temperature and relative humidity measurements were performed in the greenhouse facilities of CIRAD with two genotypes from panel B in April 2020. The sensors were placed at 15 cm above the soil surface (i.e. few centimeters above the pot brim). Recording of data started at fifteen days in the greenhouse. At 16 and 22 days after sowing, leaf area index (LAI) was measured in both low and high-density canopies using a light sensor (Spectrol LI-80, Li-Cor).

In both experiments, the sensors were covered with open polystyrene boxes to avoid direct solar radiation. Temperature and relative humidity were recorded every 30 min during a 15 days period in the glasshouse and during 6 weeks in the field trial allowing to calculate VPD.

### Measurement of LAI and light penetration in the canopy in Senegal

In the field trial at Senegal, leaf area index (LAI) was measured using PPFD measurement performed in both low and high-density canopies using a light sensor (LI-190R-BNC-2 Quantum Sensor) placed above the canopy, and a sensor bar (LI-191R-BNC-2 Line Quantum Sensor) placed at the ground level. LAI was calculated indirectly as 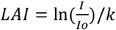 where *Io* is the incident light above the canopy, I is the light at ground level, and *k* is a crop extinction equal to 0.6 for sorghum. The Photosynthetic Photon Flux Density (PPFD) was also measured above the canopy and at two heights in the canopy on days 45 and 60 after sowing. The measurements were done with a light sensor (LI-190R-BNC-2 Quantum Sensor) placed above the canopy, and two sensor bars (LI-191R-BNC-2 Line Quantum Sensor) fixed perpendicularly on a pole placed vertically in the plots and adjusted respectively at mid-canopy level and at ground level. Mid canopy height was assessed for each genotypes thanks to graduations marks on a vertical pole. All sensors were connected to a data logger (LI-1500 Light Sensor Logger), and data were recorded between 12:30 to 1:30 pm. Each data point was the mean of four replicates in each plot. At 44 days after sowing, a light measurement was also done at ground level in a plot of high and low density at four time-point during this particular day, at 7am, 10am, 2pm and 5pm.

### Statistical analysis

Statistical analysis presented in this study were performed using GraphPad Prism version 9.2.0 for Windows, (GraphPad Software, San Diego, California USA,”http://www.graphpad.com) Slopes and time courses of transpiration were smoothed and analyzed using R software version 4.1.2. Broad sense heritability of the slopes was computed as in Falconer et al, 2005 with 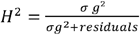 where σ p^2^ is the genotypic variance.

## Results

### The response of evapotranspiration to the evaporative demand was genotype-dependent

We examined the response of each studied genotype to the reference evapotranspiration (ET_ref_), which takes both VPD and light into account. Indeed, light intensity underwent large fluctuations in our experiments with time of day, between days and between seasons (Fig. 1), so regression lines presented in Fig. 2 correspond to evaporative demands varying with both VPD and light. Point clouds correspond to coupled values, for one genotype, of measured evapotranspiration and ET_ref_, calculated for periods of 7 to 10 days during either the dry or rainy seasons in India, or during the dry season in Senegal, at two plant densities.

**Figure 2:**
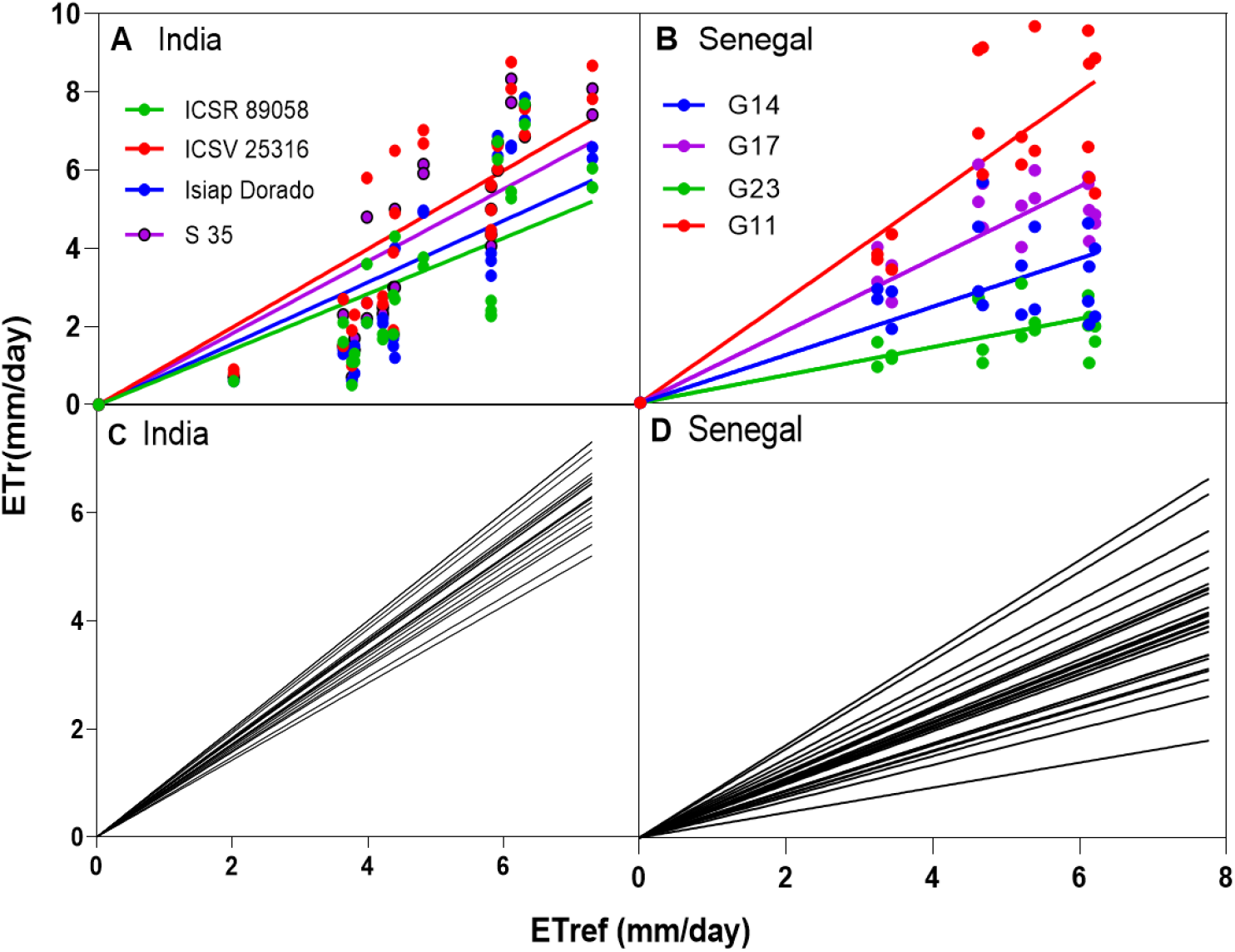
Measured evapotranspiration (ETr) plotted against the reference evapotranspiration (ETref), for 20 genotypes in the lysimeter platform in India (A, C) and 27 genotypes in the lysimeter platform in Senegal (B, D). A and B: point cloud regressions for 4 contrasting genotypes, C and D regression lines for all studied genotypes (one line per genotype). Data are means of 4 replications per genotype and treatment (P-values < 0.0001).

In the dataset collected in India, linear regressions were significant for all tested genotypes (p-values <0.0001; Suppl. table 1). We tested the linearity of relationships by considering a quadratic term in regressions, which was significantly positive for all hybrids (values of ET slightly higher than those expected under a linear assumption, Suppl. table 1, Suppl. fig 1). Hence, we never observed a tendency towards a plateau of evapotranspiration at high evaporative demands. Notably, all regressions with VPD alone were also significant, and none of them showed a tendency to plateau (non-significant quadratic term in 25% of cases, positive term otherwise). In order to facilitate genetic analyses, and in view of the small contribution of positive quadratic terms, we only considered the linear relationships in the following analyses. The slopes of linear regressions significantly differed between genotypes (Fig. 2, p-value = 0.03), with a high heritability (H^2^ = 0.56, Fig. 2 A&C). Hence, the studied genotypes differed in their ability to transpire at a given evaporative demand with up to three-fold differences, e.g. from 2.3. to 6.28 mm d^-1^ at an ET_ref_ of 5.80 mm d^-1^. Importantly, the slopes of relationships for each genotype were independent of leaf area for both the first (r = 0.2, p-value=0.42) and the second (r= 0.03, p-value=0.86) measurement in the adjacent field, so the differences between genotypes were probably not related to leaf area, and linked to stomatal behaviour (Suppl. fig 2). The same set of results was observed whether ET_ref_ was calculated either via the Penman-Monteith equation corrected for leaf area, or with the APSIM model, with a very high correlation between calculated slopes (Suppl. fig 3 A).

**Table 1:** ANOVA table showing the genotypic variability for the water use efficiency (WUE) in the dry and rainy season experiment in India and the dry season experiment in Senegal. *, p-value <0.05, **, p-value <0.01, *** p-value <0.001, ****, p-value <0.0001.

| Source of Variation | Dry season<br>2018 | Rainy season<br>2018 | Dry season<br>2021 |
| --- | --- | --- | --- |
| Location | India | India | Senegal |
| Genotypes panel | A | A | B |
| Two-way ANOVA |  |  |  |
| Genotype | * | **** | * |
| Genotype x Season | *** |  | NA |

The dataset collected in Senegal provided consistent results, with regressions between evapotranspiration and ET_ref_ significant for all studied genotypes (Fig. 2B&D, p-values <0.0001; Suppl. table 2). Quadratic terms were non-significant in 56% of cases and slightly positive otherwise (Suppl. table 2), so following analyses only considered the linear regressions as in panel A. The genetic variability of slopes was even larger than in experiments in India (p value = 0.02), with slopes ranging from 0.26 to 0.96, and a heritability of slopes of H^2^ = 0.41. For example, transpiration rate ranged from 1.03 to 9.14 mm d^-1^ at an ET_ref_ of 5.83 mm d^-1^ (Fig. 2B & D). The low range of VPD in this dataset resulted in the fact that regressions of ET with VPD alone were non-significant in most cases. Hence, regressions with ET_ref_ provided more insights than those with VPD alone in this case. Consistent results were again observed in the greenhouse, where the slopes of the transpiration response to the evaporative demand differed between genotypes (p-value < 0.01, H^2^ = 0.39) (data not shown).

### A large genotypic variability for water use efficiency

WUE showed a high genotypic variability in panel A during both the dry and rainy seasons in India (2.2 to 3.2 and 8.4 to 11.9 g kg^-1^, respectively, p-value = 0.01 and < 0.0001), with a high heritability calculated across seasons (H^2^ =0.64) (Fig. 3A and 3B). A significant genotype x season interaction was observed, suggesting that the decrease in WUE with ET_ref_ depended on other traits that differ between genotypes (p-value = 0.0004) (Table 1). Similar results were observed in Senegal with panel B, with a high genetic variability for WUE (1.4 to 3.9 g.kg^-1^, p-value < 0.01) and a high heritability (H^2^ = 0.65, Fig. 3C).

**Figure 3:**
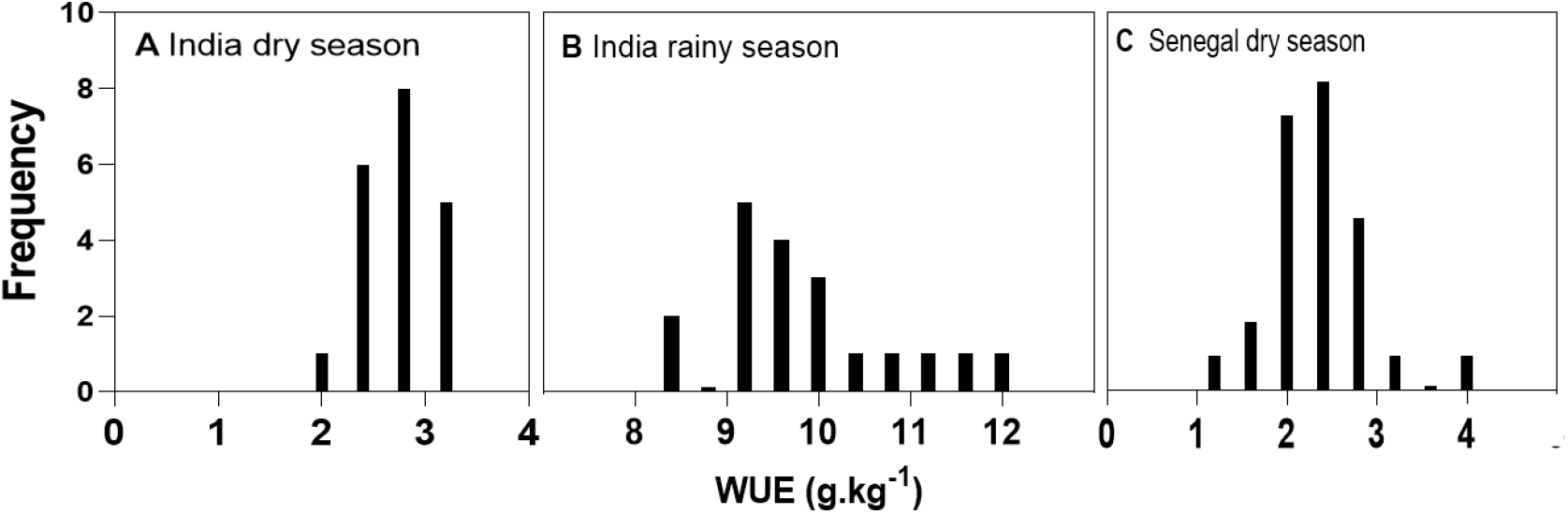
Frequency distribution of the water use efficiency (WUE) values measured on the lysimeter platforms for the 20 genotypes from panel A India during the 2018 dry (A) and rainy seasons (B), and for the 27 genotypes from panel B in Senegal during the 2021 dry season (C).

### The genetic correlation between WUE and evapotranspiration was positive under high evaporative demand, and negative under low evaporative demand

We then examined the genetic link between WUE and the response of transpiration rate to evaporative demand. In experiments performed in dry seasons of India or Senegal (Fig. 4 A and B), WUE correlated positively and significantly with the slope of the regressions presented in Fig. 2 for the same genotypes. Counter intuitively, the genotypes that most transpired at a given ET_ref_ had the highest WUE, with a high correlation between WUE and the slope of response curves in India (r = 0.79, P value <0.0001,) (Fig 4A) and lower but significant in Senegal (r = 0.54, p-value <0.01) (Fig 4B). The same pattern was observed in the greenhouse under high VPD, where the slope of the regression between transpiration and time at high transpiration hour, was positively correlated with WUE (r = 0.78, p-value < 0.05) (Suppl. fig.4). Conversely, this relationship was reversed and strongly negative in the rainy season of India (r = -0.71, p-value <0.0001, Fig 4 C). That is, the genotypes that most transpired in response to ET_ref_ had the lowest WUE in this case. This result held whether ET_ref_ was calculated via either the APSIM model or the Penman Monteith equation corrected for leaf area (Suppl. fig 3B and C).

**Figure 4:**
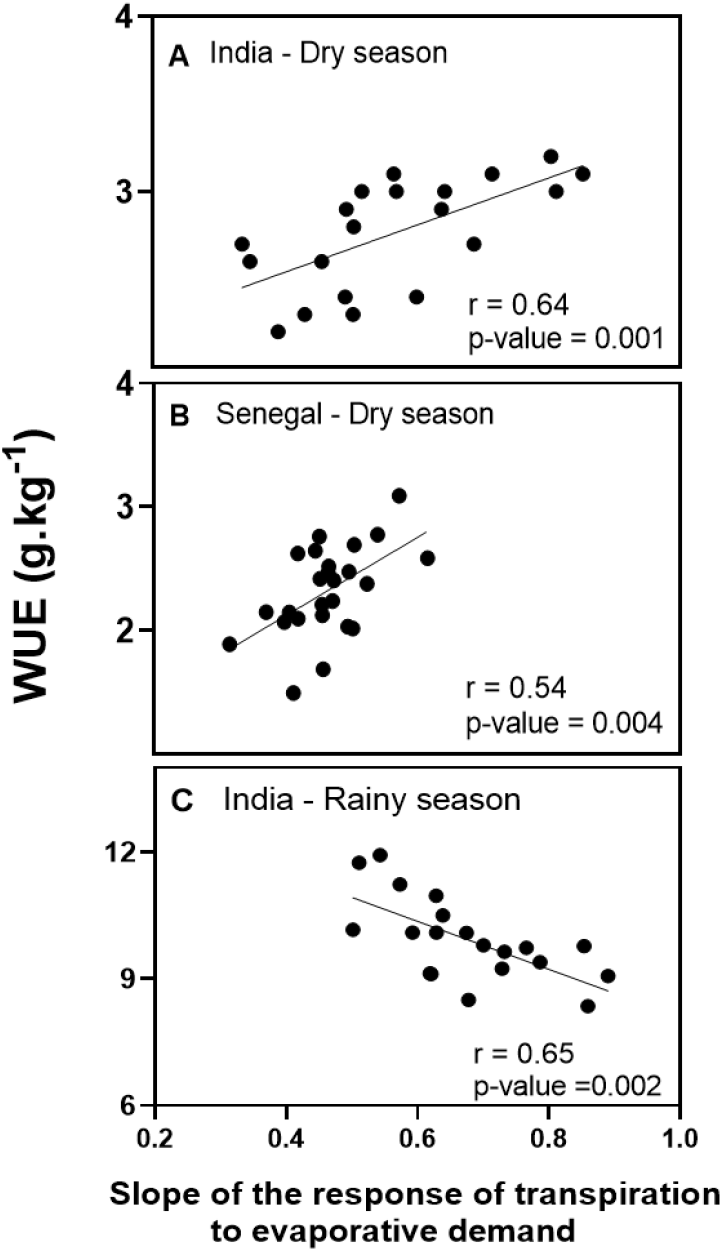
Water use efficiency (WUE) plotted against the slopes of the evapotranspiration response to ETref (see figure 2). (A) 20 genotypes of panel A in the dry season, India (r = 0.64, p-value = 0.0001). (B) 27 genotypes of panel B in the dry season, Senegal (r= 0.54, p-value = 0.004). (C) 20 genotypes of the panel A in the wet season, India (r = - 0.65, p-value < 0.01). Data are means of 4 replications per genotype and treatment (P-values < 0.0001).

### The intra canopy VPD was lower than air VPD during the day

The measured VPD was lower within the canopy than in the air, to a greater extent in high than in low plant density. In high-density canopies, this difference became significant from day 42 onwards (Fig. 5A), when LAI was approximately 2.9 (measured on day 45). In low-density canopies, it became significant 14 days later (day 56) when LAI was also close to 3 (3.2 on day 53) (Fig. 5A). Differences in VPD were maximum in the morning and early afternoon (average difference of 0.97 kPa, p-value = 0.05 over this period, Fig 5B). Differences in VPD between canopy and air were also observed in the greenhouse experiment during a 15-days period, with a mean difference in VPD of 0.63 kPa (0.55 to 0.93 kPa, p-value = 0.001, Suppl. fig 5). Overall, VPD within the canopy was lower than that sensed by upper leaves in direct relation to open air, and more so in dense canopies. This was likely due to the intra canopy transpiration because differences nullified during the night and increased with LAI.

**Figure 5:**
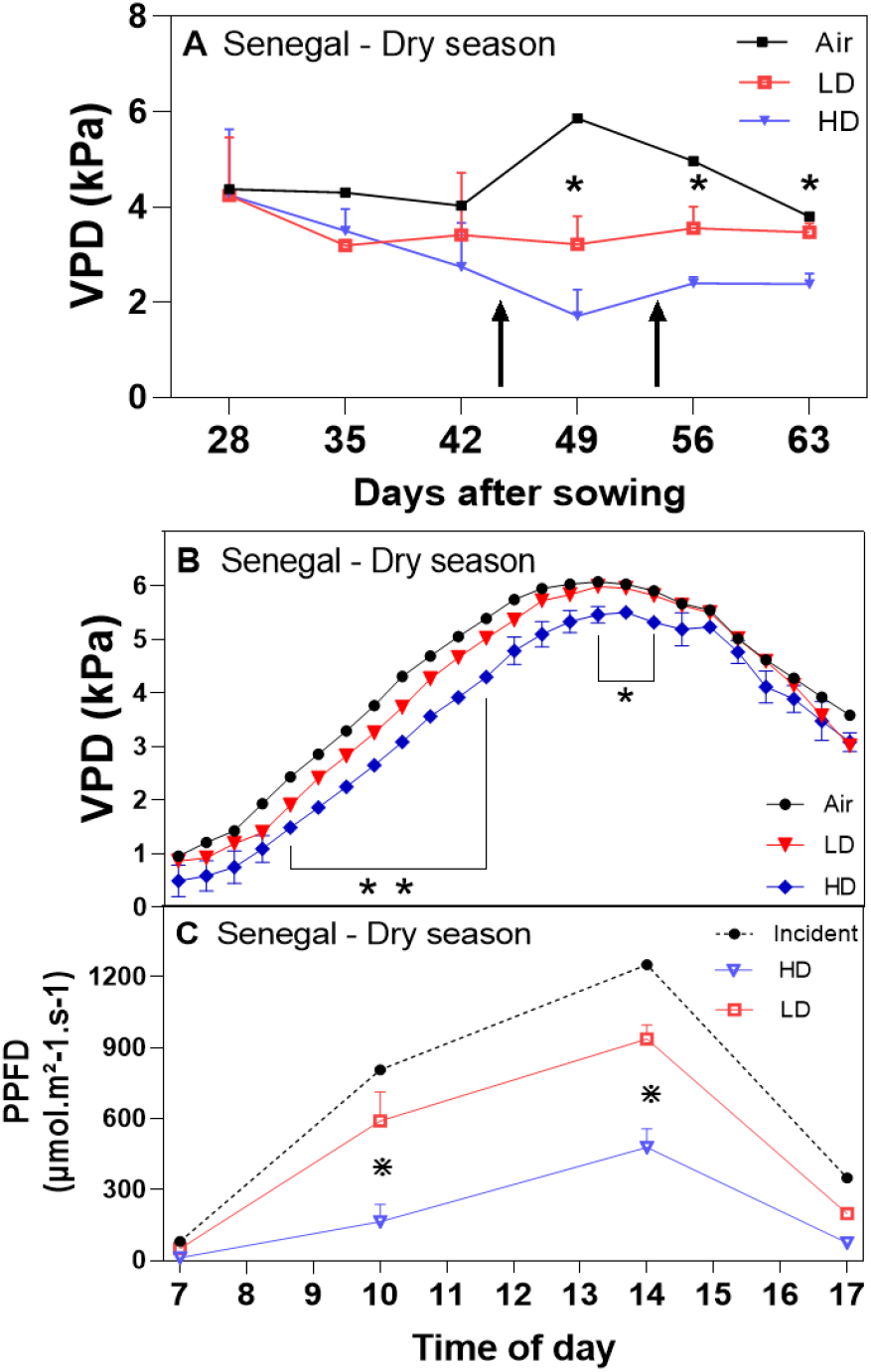
(A) Vapor pressure deficit (VPD) measured in the air and within canopies with high and low densities (12 and 24 plants/m^2^ respectively, HD and LD), as a function of time after sowing in the dry season field trial in Senegal. Black arrows in (A) represent dates at which LAI was measured (45 and 53 days after sowing), and stars indicates significant differences between HD and LD (paired t-test, p-value <0.001). (B) Daily time course of VPD during the 4th week of the same field experiment from 7am to 7 pm, where stars indicate significant differences between HD and LD (paired t-test, p-value <0.001). (C) Photosynthetically active Photon Flux Density (PPFD) at 4 time points across the same day in the two densities. Stars indicates significant differences between HD and LD (paired t-test, p-value <0.0001). Each data point is the average of sensor data collected in three plots for each of the densities.

### Is the positive relation between WUE and transpiration linked to the vertical distribution of VPD and light penetration in the canopy?

We measured light penetration in the canopy to better understand the positive genetic correlation between WUE and high transpiration under high VPD. In the experiment in India, light measurement on day 44 showed that light was appreciable within the canopy, to a larger extent for canopies with low than high density, at 10am (488 vs 165 μmol m^-2^ s^-1^ respectively) and 2pm (958 vs 479 μmol m^-2^ s^-1^ respectively) (Fig 5 C).

In the Senegal experiment, genotypes that most let the light penetrate at mid-canopy level were those that had highest WUE at 60 DAS, when LAI was 3.9 (r = 0.44, p-value = 0.009) (Fig. 6B). At this phenological stage, the genotypic correlation was observed with light intensity at mid-canopy level and not at ground level because studied canopies intercepted nearly all the incident light. At an earlier stage (45 DAS, LAI = 2.9), the genotypic correlation was observed with light at ground level, reached by light at this stage (r = 0.56, p-value = 0.04) (Fig. 6A). Hence, genotypes that most transpired at a given evaporative demand were also those for which more light reached leaves inside the canopy, potentially increasing the photosynthesis of these leaves. Because the intra-canopy VPD was lower than air VPD (Fig. 5A&B), the response of transpiration rate to ET_ref_ had a higher effect on biomass than on transpiration, thereby increasing WUE (Suppl. fig. 6A). This was not observed in the rainy season (Suppl. fig. 6B), probably explaining the observed negative genotypic effect of transpiration on WUE. Hence, we raise the possibility that genetic correlations between transpiration and WUE depend on plant architecture and the vertical distribution of VPD, with a positive relationship due to light penetration under the high light and VPD observed during dry seasons, and a negative relationship under lower light and VPD observed during the studied rainy season.

**Figure 6:**
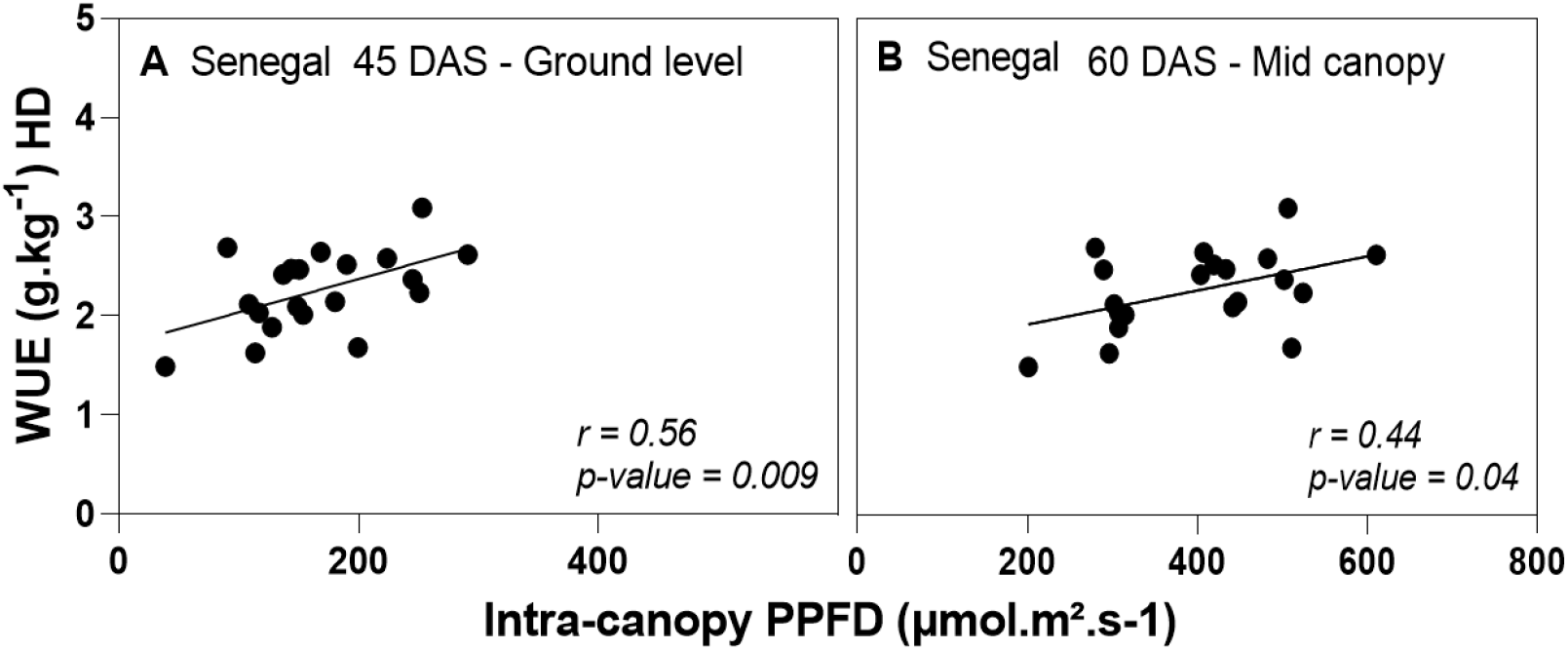
Water use efficiency (WUE) measured in lysimeters plotted against the amount of light measured at two altitude in the canopy. Senegal dry season, at (A) 45 DAS (ground level) and (B) 60 DAS (mid-canopy level). Data are the mean of four replications for WUE and three replications for light measurements.

## Discussion

### A large genetic variability of the response of transpiration to evaporative demand in natural conditions

This work extends to natural conditions, with high light and VPD, the observation of a genetic variability of the response of transpiration to evaporative demand, of particular importance for dry tropical areas. Here, we considered evaporative demand via its two components, light intensity and VPD, and observed significant regressions between the resulting ETref and evapotranspiration of each of the 47 tested genotypes. Because slopes significantly differed between genotypes in both panels A and B, we can conclude that a significant genetic variability exists for the ability of a genotype to transpire at a given evaporative demand. This variation was not related to differences in leaf area. Given the high heritability of slopes, this genetic variability may be used for designing genotypes for either high transpiration rate (suited to favourable environmental scenarios) or lower transpiration rate (suited to drought-prone areas). Notably, the accessions of panel A had a lower genetic variability and heritability for slopes than the elite hybrids of panel B, thereby suggesting that the response of transpiration to evaporative demand may have faced a selection pressure. Accessions from panel A came from the ICRISAT breeding program and were mostly bred for the rainy season, which could have explained the lower variation for a trait addressing drought-prone situations.

It was considered by Sinclair and other authors that the differential responses to evaporative demand is both characterized by stomatal closure which occurs at differently high VPD for each genotype (VPD breakpoints) and by differences in the slopes before and after the breakpoint, thereby resulting in a genetic variability of responses (e.g. Fletcher et al., 2008; Kholova et al., 2010). Here, we did not observe such a non-linear response, as quadratic terms of regressions, all positive when significant, indicated a concave relationship rather than a plateau. Indeed several studies carried in the field showed responses that did not always present a non-linear response, even when the evaporative demand was expressed via VPD only (Tharanya *et al*., 2018; Devi and Reddy, 2018). In the same way, when a confusion of effect between temperature and VPD was avoided, only a small proportion of wheat genotypes presented a plateau under high evaporative demand (Tamang *et al*., 2022). Importantly, some of the genotypes that did not present a plateau in our study, did show a marked plateau at high VPD in a study in growth chamber (Karthika *et al*., 2019). Hence, a different representation emerges from studies carried out in high evaporative demand carried out outdoor compared with those in growth chamber. Because no plateau for evaporative demand was observed, the cause of the genetic variability of the evapotranspiration was not stomatal closure at high VPD, but rather a difference in transpiration rate over the whole range of evaporative demand. Indeed, an appreciable genetic variability for stomatal conductance was observed in maize, even at relatively low evaporative demands, with high heritability and consistent QTLs (Alvarez Prado et al 2019, Welcker et al 2022). We therefore suggest that genetic differences in transpiration rate were due to intrinsic differences in stomatal conductance, rather than to an adaptive process of stomatal closure at high VPD.

Why different shapes of the response of transpiration to evaporative demand were observed between results in growth chamber and in our field study? We raise the possibility that this is due to the difference in light intensities between the two conditions. Under relatively low light and high air movement, as in a growth chamber, the decrease in stomatal conductance with high VPD directly translates into a reduction in transpiration rate. In conditions with high light and lower wind, uncoupling occurs between stomatal conductance and transpiration for most species (Jarvis and Mcnaughton, 1986), so leaf temperature increases with stomatal closure, thereby increasing leaf-to-air vapour pressure difference, largely decreasing the effect of stomatal closure on transpiration rate (Chaves *et al*., 2016). It was unexpected that a good relationship between transpiration and air VPD was still observed in our experiments in India, as it did in former studies (Kar et al., 2020; Kholova et al., 2010). This was probably due to the correlation between light intensity and VPD, observed in our study, but probably not in other climatic conditions, for example in regions where the wind brings either dry air from continental areas, or wet air from the sea, depending on its direction, with an unchanged light intensity (Ben Haj Salah and Tardieu, 1996). Indeed, this correlation was not observed in the Senegal experiment.

### A genetic variability of WUE related to the response of evapotranspiration to evaporative demand

The results presented here suggest that lower VPD within the canopy may increase the WUE of leaves transpiring within the canopy, during seasons facing high VPD, because WUE is inversely related to VPD (Sinclair, Tanner and Bennett, 1984; Condon *et al*., 2002). We propose that observed differences in WUE among genotypes were, in part, a consequence of the proportion of transpiration contributed by leaves within the canopy. The large and heritable genotypic variation response of evapotranspiration to ET_ref_ supports this hypothesis. Our interpretation is that a stronger evapotranspiration response would come from more leaves actively participating in plant transpiration, i.e. by involving leaves inside the canopy. Genotypes with highest WUE in high VPD seasons were those for which more light was available inside the canopy. Architectural features could be involved in the variability of response, with genotypes that allow the lower canopy levels to participate in transpiration, benefitting from lower VPD values, allowing for an increase in WUE over the long term. Indeed plant architecture strongly influences variables such as light interception (Falster and Westoby, 2003; Duursma *et al*., 2012; Iii *et al*., 2015) and radiation use efficiency (RUE) (George-jaeggli *et al*., 2013; Truong *et al*., 2015; Perez *et al*., 2019). Other studies have also shown a better efficiency of water use in varieties allowing a better distribution of the light resource in other species (Falster and Westoby, 2003; Lee and Tollenaar, 2007).

### Why WUE was highest in genotypes with high response to transpiration in dry seasons and lowest in wet season?

WUE was either positively or negatively genetically correlated to the evapotranspiration response to the evaporative demand, depending on the prevailing light and VPD during the season. This contrasts with earlier results showing that it is related to transpiration restriction under high VPD (Sinclair, Hammer and Van Oosterom, 2005; Vadez *et al*., 2014). We propose that differences in WUE may be related to differences in the distribution of the light resource inside the canopy, not taken into account in earlier studies that only considered VPD for variation of transpiration. During the dry season with high VPD and light, highly responsive genotypes for transpiration also allowed light to penetrate deeper in the canopy, letting lower level leaves to transpire and photosynthesize. This caused proportionally more transpiration to occur for intra-canopy than for top leaves in genotypes with high transpiration. Intra-canopy leaves benefited from the low VPD and they could maintain photosynthesis better than genotypes with lower light penetration, and at a lower water cost than genotypes photosynthesizing mostly from the top leaves. Accordingly, the increase in biomass was proportionally higher than that of the evapotranspiration in the high VPD season (Suppl fig. 3A), suggesting a positive trade-off of biomass for water.

On the contrary, during the rainy season with less incident light and VPD, a higher slope of the ETr response to ETref correlated to a lower WUE. Our interpretation is that light resource was insufficient to penetrate deeply inside the canopy, so both transpiration and light interception essentially involved the top leaves in the canopy, i.e. those exposed more to air VPD of the environment, VPD value that were still above 2kPa during that season. Hence, we propose here that differences in canopy architecture allowed variations in the light available inside the canopy, which drove transpiration under high evaporative demand, and increased WUE because of a higher proportion of transpiration benefitting from milder VPD conditions allowed by dense canopies.

## Acknowledgement

The senior and corresponding authors are in part supported by a grant the Make Our Planet Great Again (MOPGA) ICARUS project (Improve Crops in Arid Regions and Future Climates) funded by the Agence Nationale de la Recherche (ANR, grant ANR-17-MPGA-0011).

## Supplementary figure and tables captions

*Suppl. Fig 1:* Example of the polynomial regressions using both VPD (grey lines) and ETref (red lines) in x axis for two genotypes from the panel A. Genotypes MR750 showed significant quadratic term with both variables with a concave relationship (A). Genotype NTJ-2 showed non-significant quadratic term with VPD and significant one with concave relationship with ET ref.

*Suppl. Fig 2:* Leaf area index (LAI) measured 33 (A) and 40 (B) days after sowing in the 2018 dry season field experiment as a function of the slope of the transpiration response to evaporative demand (ETref) from the adjacent lysimetric experiment. (India, Panel A).

*Suppl. Fig 3:* Linear regression of the slopes values generated with Kc method as a function of the values of the slopes generated with APSIM for the dry season experiment in India (r= 0.99) (A). Water use efficiency (WUE) as a function of the slope generated by the regression of the measured evapotranspiration against the ETref calculated with the Penman-Monteith corrected with a crop constant (Kc) for the 20 genotypes of the panel A in the dry (B) and rainy (C) season in India.

*Suppl. Fig 4:* Water use efficiency (WUE) plotted against the slope of the time course of transpiration rate during the 3 hours preceding the maximum transpiration in the indoor lysimeter experiment (Montpellier, France, 9 genotypes from panel B).

*Suppl. Fig 5:* Vapor pressure deficit (VPD) measured in air and within canopies with high and low densities (12 and 24 plants/m^2^ respectively, HD and LD), as a function of time after sowing in the glasshouse experiment (Montpellier, France, 2 genotypes from panel B). Stars indicates significant differences between air and HD VPD.

*Suppl. Fig 6:* Normalized biomass and evapotranspiration (ET) plotted against the slopes generated by the regression of measured evapotranspiration and ET_ref_ during the dry (A) and wet (B) seasons in India.

**Suppl. table 1:** Values of slopes, R^2^, significance of the regressions, quadratic terms of the regressions and significance of the quadratic terms for all genotypes from the panel A tested in India. Regressions are ETr values from the lysimeters as a function both ETref (calculated via APSIM software) or VPD (Calculated with T° et RH data from the adjacent meteorological station). *, p-value <0.05, **, p-value <0.01, *** p-value <0.001, ****, p-value <0.0001.

| Genotypes<br>Panel A | Slopes<br>($ET_{ref}$) | $R^2$<br>($ET_{ref}$) | Slopes<br>p-values | $x^2$<br>term | $X^2$<br>term p-value | Genotypes<br>Panel A | Slopes<br>(VPD) | $R^2$<br>(VPD) | p-values | $x^2$<br>term | $X^2$ term<br>p-value |
| --- | --- | --- | --- | --- | --- | --- | --- | --- | --- | --- | --- |
| CSH 16 | 0,86 | 0,65 | <0,0001 | 0,09 | 0,021 | CSH 16 | 1,20 | 0,62 | <0,0001 | 0,08 | 0,150 |
| ICSB 404 | 0,74 | 0,65 | <0,0001 | 0,14 | 0,005 | ICSB 404 | 1,04 | 0,68 | <0,0001 | 0,23 | 0,000 |
| ICSH<br>14002 | 0,91 | 0,75 | <0,0001 | 0,13 | 0,408 | ICSH<br>14002 | 1,27 | 0,74 | <0,0001 | 0,15 | 0,079 |
| ICSH<br>28001 | 0,96 | 0,75 | <0,0001 | 0,13 | 0,322 | ICSH<br>28001 | 1,34 | 0,73 | <0,0001 | 0,12 | 0,074 |
| ICSR 101 | 0,80 | 0,73 | <0,0001 | 0,11 | 0,067 | ICSR 101 | 1,12 | 0,74 | <0,0001 | 0,14 | 0,004 |
| ICSR<br>14001 | 0,85 | 0,75 | <0,0001 | 0,15 | 0,096 | ICSR<br>14001 | 1,19 | 0,73 | <0,0001 | 0,17 | 0,003 |
| ICSR 196 | 0,86 | 0,73 | <0,0001 | 0,15 | 0,305 | ICSR 196 | 1,21 | 0,74 | <0,0001 | 0,18 | 0,001 |
| ICSR<br>89058 | 0,71 | 0,61 | <0,0001 | 0,12 | 0,010 | ICSR<br>89058 | 1,00 | 0,64 | <0,0001 | 0,20 | 0,000 |
| ICSV 112 | 0,84 | 0,76 | <0,0001 | 0,17 | 0,236 | ICSV 112 | 1,17 | 0,75 | <0,0001 | 0,21 | 0,001 |
| ICSV<br>15013 | 0,91 | 0,70 | <0,0001 | 0,15 | 0,282 | ICSV<br>15013 | 1,27 | 0,67 | <0,0001 | 0,14 | 0,036 |
| ICSV<br>25302 | 0,98 | 0,73 | <0,0001 | 0,12 | 0,455 | ICSV<br>25302 | 1,36 | 0,67 | <0,0001 | 0,10 | 0,349 |
| ICSV<br>25308 | 1,00 | 0,69 | <0,0001 | 0,10 | 0,371 | ICSV<br>25308 | 1,39 | 0,62 | <0,0001 | 0,05 | 0,452 |
| ICSV<br>25316 | 1,00 | 0,71 | <0,0001 | 0,10 | 0,354 | ICSV<br>25316 | 1,39 | 0,65 | <0,0001 | 0,06 | 0,330 |
| ICSV 745 | 0,90 | 0,74 | <0,0001 | 0,18 | 0,236 | ICSV 745 | 1,26 | 0,76 | <0,0001 | 0,22 | <0,0001 |
| ICSV<br>93046 | 0,86 | 0,69 | <0,0001 | 0,16 | 0,145 | ICSV<br>93046 | 1,20 | 0,65 | <0,0001 | 0,18 | 0,043 |
| Isiap<br>Dorado | 0,79 | 0,71 | <0,0001 | 0,16 | 0,005 | Isiap<br>Dorado | 1,11 | 0,77 | <0,0001 | 0,24 | <0,0001 |
| MR 750 | 0,82 | 0,73 | <0,0001 | 0,18 | 0,002 | MR 750 | 1,15 | 0,79 | <0,0001 | 0,26 | <0,0001 |
| NTJ-2 | 0,91 | 0,70 | <0,0001 | 0,13 | 0,368 | NTJ-2 | 1,26 | 0,67 | <0,0001 | 0,12 | 0,230 |
| PVK 801 | 0,89 | 0,74 | <0,0001 | 0,15 | 0,053 | PVK 801 | 1,26 | 0,77 | <0,0001 | 0,20 | 0,000 |
| S 35 | 0,92 | 0,75 | <0,0001 | 0,15 | 0,319 | S 35 | 1,29 | 0,74 | <0,0001 | 0,14 | 0,017 |
| p-values | 0,03 |  |  |  |  | p-values | 0,0003 |  |  |  |  |
| H<sup>2</sup> | 0, 56 |  |  |  |  | H<sup>2</sup> | 0,66 |  |  |  |  |

**Suppl. table2:** Values of slopes, R^2^, significance of the regressions, quadratic terms of the regressions and significance of the quadratic terms in all genotypes from the panel B. Regressions are ETr values from the lysimeters as a function of Etref calculated via the Penman-Monteith equation corrected with a crop constant (Kc). *, p-value <0.05, **, p-value <0.01, *** p-value <0.001, ****, p-value <0.0001.

| Genotypes<br>Panel B | Slopes<br>(E<sub>Tref</sub>) | R<sup>2</sup> (E<sub>Tref</sub>) | Slopes p-values | x<sup>2</sup> term | X<sup>2</sup> term p-value |
| --- | --- | --- | --- | --- | --- |
| *IS 11119* | 0,85 | 0,61 | <0,0001 | 0,15 | 0,010 |
| *CE-145-66* | 0,91 | 0,35 | <0,0001 | 0,10 | 0,072 |
| *IS 3507* | 1,07 | 0,61 | <0,0001 | 0,18 | 0,023 |
| *IS 33209* | 0,73 | 0,50 | <0,0001 | 0,11 | 0,017 |
| *ICSV 745* | 0,63 | 0,59 | <0,0001 | 0,09 | 0,186 |
| *IS 14539* | 0,50 | 0,25 | <0,0001 | 0,04 | 0,512 |
| *IS 33173* | 0,74 | 0,32 | <0,0001 | 0,10 | 0,079 |
| *IS 13452* | 0,68 | 0,57 | <0,0001 | 0,10 | 0,018 |
| *FAURO* | 0,74 | 0,66 | <0,0001 | 0,11 | 0,079 |
| *IS 19016* | 0,64 | 0,27 | <0,0001 | 0,10 | 0,166 |
| *IS 16396* | 0,75 | 0,43 | <0,0001 | 0,11 | 0,014 |
| *IS 7958* | 0,64 | 0,37 | <0,0001 | 0,16 | 0,005 |
| *IS 14556* | 0,50 | 0,65 | <0,0001 | 0,04 | 0,346 |
| *IS 15443* | 0,64 | 0,31 | <0,0001 | 0,07 | 0,274 |
| *IS 14963* | 0,67 | 0,42 | <0,0001 | 0,08 | 0,075 |
| *IS 13845* | 0,29 | 0,39 | <0,0001 | 0,02 | 0,319 |
| *IS 2678* | 0,54 | 0,51 | <0,0001 | 0,06 | 0,030 |
| *IS 25207* | 0,53 | 0,47 | <0,0001 | 0,06 | 0,266 |
| *E36-1* | 0,54 | 0,21 | <0,0001 | 0,07 | 0,320 |
| *IS 31202* | 0,80 | 0,75 | <0,0001 | 0,12 | 0,037 |
| *IS 15702* | 0,47 | 0,58 | <0,0001 | 0,07 | 0,252 |
| *IS 33423* | 0,42 | 0,50 | <0,0001 | 0,06 | 0,271 |
| *IS 33261* | 0,66 | 0,47 | <0,0001 | 0,09 | 0,070 |
| *IS 16125* | 0,66 | 0,39 | <0,0001 | 0,12 | 0,068 |
| *IS 2986* | 1,02 | 0,44 | <0,0001 | 0,11 | 0,027 |
| *IS 10876* | 0,61 | 0,63 | <0,0001 | 0,10 | 0,070 |
| *PAYENNE* | 0,63 | 0,71 | <0,0001 | 0,10 | 0,014 |
| p-values | 0,02 |  |  |  |  |
| H<sup>2</sup> | 0,41 |  |  |  |  |

